# Subcellular dynamics of leghemoglobin is modulated by its site-specific serine phosphorylation during symbiotic nitrogen fixation in *Lotus japonicus*

**DOI:** 10.64898/2026.04.23.719809

**Authors:** Md Azimuddin Ashrafi, Aniruddho Das, Anirban Siddhanta

## Abstract

Symbiotic nitrogen fixation (SNF) relies on aerobic respiration, yet the key enzyme, nitrogenase, is extremely oxygen labile. Leghemoglobin (Lb) resolves this “oxygen paradox” by buffering and facilitating O_2_ transport. However, the dynamic regulation of Lb during nodule development remains poorly understood. Earlier results from our laboratory demonstrated that site-specific serine phosphorylation of Lb reduces its oxygen sequestration capacity. Here, we investigated the spatio-temporal regulation of Lb with the progress of rhizobial load during SNF. Fluorescence immunohistochemistry (FIHC) using anti-Lb antibody revealed that its localization gradually shifted from the plasma membrane to the cytoplasm of infected cells as nodules mature. Using phospho-peptide (Lb) specific antibodies, we found that serine phosphorylation triggers this translocation. Furthermore, FIHC in conjunction with immunoprecipitation followed by immunoblotting with phospho- and non-phospho-peptide specific antibodies demonstrated that the non-phosphorylated form is detectable as early as 9 dpi, whereas the phosphorylated forms were first detected at 11 dpi and progressively accumulated during nodule maturation. This spatio-temporal transition coincides with increasing rhizobial colonization and is accompanied by a decline in the non-phosphorylated pool. Therefore, the increased cytoplasmic pool of phosphorylated Lb, which exhibits reduced oxygen sequestration capacity, likely functions in promoting oxygen transport to sustain elevated rhizobial respiration. Together, these findings demonstrate that site-specific serine phosphorylation represents one of the key regulatory mechanisms linking Lb localization dynamics with progression of rhizobial infection, thereby contributing to the maintenance of oxygen homeostasis during SNF.

## Introduction

The symbiotic relationship between leguminous plants and nitrogen-fixing rhizobia is a cornerstone of sustainable agriculture, culminating in the formation of root nodules where atmospheric dinitrogen (N_2_) is converted to ammonia. This process is energetically expensive and requires a high flux of oxygen for aerobic respiration (Appleby, 1984; Udvardi and Poole, 2013). In contrast, the nitrogenase enzyme complex that catalyzes ammonia formation is irreversibly inactivated by oxygen. This “oxygen paradox” is primarily managed by leghemoglobins (Lbs), specialized plant hemeproteins that accumulate millimolar concentrations in infected nodule cells (Appleby, 1984; Ott et al., 2005).

Lbs exhibit high O_2_ affinity, enabling efficient oxygen delivery to respiring rhizobia while maintaining the low free-O_2_ environment required for nitrogenase activity. (Appleby, 1984; Udvardi and Poole, 2013). Despite the well-established biochemical properties of Lb, the mechanisms that dynamically regulate its activity *in vivo* to accommodate changing metabolic states of the developing nodule is not fully understood.

Previous work from our laboratory showed that *in vitro* phosphorylation of Lb from *L. japonicus* at a specific serine residue (S45) significantly reduces its oxygen sequestration capacity by disrupting the heme-binding pocket (Bhar et al., 2015). Our finding was validated by phosphoproteomic studies in *Medicago truncatula* that identified phosphorylation at a homologous site, suggesting a conserved regulatory role (Marx et al., 2016; Lin, 2018). Besides phosphorylation, interaction with other nodule proteins plays important role in the regulation of Lb activity. Indeed, we previously demonstrated that Lb interacts with Nodulin 16 (Nlj16), another late nodulin in *L. japonicus* (Ghosh et al., 2019). This interaction enhances Lb’s oxygen-binding affinity, an effect opposite to that of serine phosphorylation. Furthermore, our preliminary observation strongly indicates that Nlj16 is essential for recruiting Lb to the cell membrane of infected cells (Ghosh et al., 2022).

This suggests a sophisticated regulatory system where the balance between Lb phosphorylation and its interaction with Nlj16 could fine-tune oxygen homeostasis in a determinant nodule during SNF. However, the spatial and temporal dynamics of these regulatory events within the context of a developing nodule have not been explored.

In this study, we investigated the in vivo spatiotemporal dynamics of Lb and its phosphorylation in determinate nodules of *Lotus japonicus* using phospho-specific peptide antibodies. We demonstrate that phosphorylation of Lb is developmentally regulated and correlates with its redistribution from the plasma membrane to the cytoplasm during nodule maturation. This transition coincides with increasing rhizobial colonization and metabolic demand, consistent with a functional shift from oxygen sequestration to controlled oxygen delivery. Together, our findings provide new insight into the molecular regulation of oxygen homeostasis in determinate nodules and identify phosphorylation-dependent modulation of leghemoglobin as a key mechanism supporting efficient symbiotic nitrogen fixation.

## Materials and Methods

### Plant Material and Growth Conditions

*Lotus japonicus* (ecotype Gifu B-129) seeds were obtained from the National Bioresource Project (University of Miyazaki, Japan). Plants were grown under controlled environmental conditions with a 16-h-light/8-h-dark photoperiod, day/night temperatures of 22°C/18°C, and 70% relative humidity. Seeds were scarified in concentrated H_2_SO_4_ for 10 min, followed by surface sterilization in 25% (v/v) commercial bleach (1% sodium hypochlorite) containing 0.1% (v/v) Triton X-100 for 10 min. Seeds were rinsed six times with sterile distilled water, soaked overnight at room temperature, and stratified at 4°C for 24 h. Seeds were germinated on 1% (w/v) agar plates containing one-quarter-strength B&D (Broughton and Dilworth 1971) medium. Seedlings were inoculated with *Mesorhizobium loti* strain NZP2235 and grown in B&D nutrient solution. For nodulation assays, plants were maintained under nitrogen-free conditions.

### Rhizobial Strain, Growth Conditions, and Inoculation

*Mesorhizobium loti* wild-type strain NZP2235 (Jarvis et al., 1982) was used for nodulation of *L. japonicus* (Gifu B-129). Rhizobial cultures were grown in yeast mannitol broth containing 0.1 g yeast extract, 0.05 g K_2_HPO_4_, 0.02 g MgSO_4_, 0.01 g NaCl, and 1 g mannitol per 100 mL medium, supplemented with 10% (v/v) of 100 mM CaCl_2_, and adjusted to pH 6.8. Cultures were incubated at 28°C in the dark for 24 h. For inoculation, plants were flood-inoculated with *M. loti* suspension at an OD_600_ of 0.03–0.04.

### Antibodies

Customized affinity purified antisera against phospho- and non phospho-peptides of leghemoglobin (Lb) containing serine residues S45 and S55 were generated in rabbit (Biotech Desk Pvt. Ltd). Customized antibodies against *Lotus japonicus* plasma membrane intrinsic protein 2 (*Lj*PIP2) were raised in rat (ABGENEX INDIA PVT.LTD). Anti-leghemoglobin antibody was kindly provided by C. P. Vance (University of Minnesota, St. Paul, USA). All anti-peptide antibodies were affinity purified using peptide-conjugated columns and validated by enzyme-linked immunosorbent assay (ELISA) and immunoblot analysis.

### Fluorescence Immunohistochemistry

FIHC was performed as described by Ghosh et al. (2019). Briefly, nodules were collected at 7– 21 dpi and embedded in Technovit 7100 resin (Heraeus Kulzer). Microtome sections (10–12 μm) were prepared and blocked in immune buffer (20 mM Tris-HCl, pH 8.2, 0.9% NaCl, 0.01% BSA, 0.02% gelatin). Sections were incubated with primary antibodies (1:100 dilution), followed by respective Alexa Fluor conjugated secondary antibodies (1:250 dilution; Invitrogen, Thermo Fisher Scientific). Confocal images were acquired using an Olympus confocal microscope (FV3000) and processed with FLUOVIEW 3000 and ImageJ software. At least 10 nodules from three separate plants were analyzed for each time point.

### Nodule Protein Extraction, immunoprecipitation and immunoblotting

Nodules were harvested at different days post-inoculation (dpi) and immediately frozen in liquid nitrogen. Samples were ground to a fine powder and homogenized in extraction buffer containing 50 mM Tris-HCl (pH 7.6), 10 mM EDTA (pH 8.0), 50 mM NaCl, 10% (v/v) glycerol, 25 mM NaF, 50 mM sodium pyrophosphate, 1 mM sodium orthovanadate, and protease inhibitor cocktail. Polyvinylpyrrolidone (2% v/v) was added to remove phenolic compounds. Homogenates were vortexed and incubated on ice for 5 min, followed by clarification at 400 × g for 10 min at 4°C. Soluble proteins were separated from bacteroids by centrifugation at 13,000 × g for 20 min at 4°C, and the supernatant was collected for further analysis.

Immunoprecipitation was performed as described by Chakrabarti et al. (2015). Briefly, 5 μg of rabbit anti-leghemoglobin (Lb) antibody was incubated with native protein extracts. Immunocomplexes were resolved by 12% SDS-PAGE, transferred to PVDF membranes, and probed with rabbit anti-phospho-(S45,55) peptide antibody, followed by HRP-conjugated secondary antibody. Signals were detected using SuperSignal^**™**^ West Pico PLUS Chemiluminescent substrate (Thermo scientific) and visualized using a ChemiDoc^™^ imaging system (BIO-RAD).

### Analysis of Rhizobial Load

Nodule sections at different dpi were stained with propidium iodide and examined microscopically. Rhizobial load was further quantified by real-time PCR analysis of the nitrogenase (*nifH*) gene transcript.

### RNA Isolation and cDNA Synthesis

Total RNA was isolated from *L. japonicus* nodules harvested at 0, 9, 14, and 21 dpi using the NucleoSpin RNA Plant Kit (Macherey-Nagel). First-strand cDNA was synthesized from 1 μg of total RNA using random hexamer primers and the SuperScript III First-Strand Synthesis System (Invitrogen).

### Quantitative Real-Time PCR

Quantitative real-time PCR (qRT-PCR) was performed using 5 ng of cDNA with the DyNAmoColorFlash SYBR Green qPCR Kit (Thermo Scientific). Gene-specific primers targeting *nifH* were used, with *sigA* serving as the internal control. Reactions were performed on an ABI 7500 Fast Real-Time PCR system. Thermal cycling conditions were as follows: 95°C for 5 min, followed by 40 cycles of 95°C for 15 s, 60°C for 30 s, and 72°C for 30 s. Relative gene expression was calculated using the 2^^−ΔΔCt^ method. Experiments included three biological replicates, each with technical replicates.

### Statistical Analysis

Data is presented as means ± SD. Statistical significance between infected samples and uninfected controls (0 dpi) was determined using paired *t*-tests (GraphPad). Significance levels were defined as *P*< 0.05, *< 0*.*01and < 0*.*001*.

## Results

### Spatio-temporal distribution of Leghemoglobin during nodule development

To investigate the developmental dynamics of Lb localization during nodulation, we performed fluorescence immunohistochemistry (FIHC) on nodules harvested at 9, 14, and 21 dpi.

At 9 dpi, Lb signal was predominantly enriched at the periphery of infected nodule cells, adjacent to the cell wall (Fig. 1A, a; triangle). By 14 dpi, Lb distribution expanded into the cytoplasm (Fig. 1A, d: star), and at 21 dpi, the protein was predominantly distributed throughout the cytoplasm (Fig. 1A, g: star). Quantitative fluorescence intensity profiles along the yellow transect lines across infected cells (Fig. 1A insets in a, d & g) confirmed this transition, revealing a shift from peripheral enrichment at early stages to a more uniform cytoplasmic distribution in mature nodules (Fig. 1B, a, d & g).

**Figure 1.**
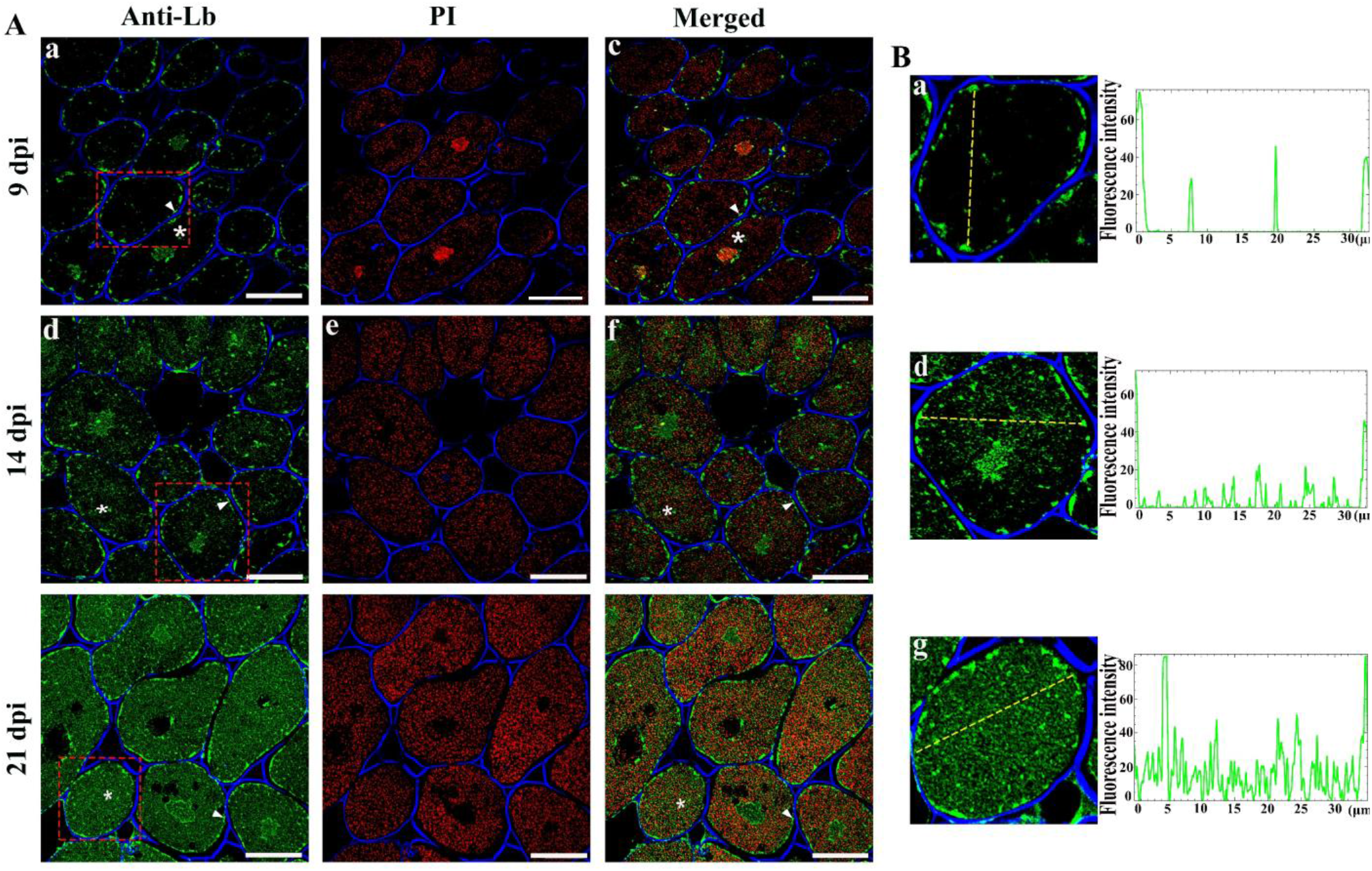
Spatio-temporal distribution of Lb in *Lotus japonicus* nodules. **(A)** Microtome sections (a–i) of nodules harvested at 9 dpi (a–c), 14 dpi (d–f) and 21 dpi (g–i) were processed for immunohistochemistry as described in Methods. Lb was detected using anti-Lb antibody (green; a, d & g). Plant and bacterial nuclei were counterstained with propidium iodide (PI; red; b, e & h). Merged images are shown in c, f and i. Cell walls were visualized with Calcofluor White (blue) in all samples. _*_ Indicates cytoplasmic and ▲demarcate cell periphery localization of Lb. Bars = 20 μm. **(B)** Fluorescence intensity profiles of Lb (green) within a nodule cell marked by the red dashed box (Panel A; a, d & g) were quantified using ImageJ. For that, three transect lines (indicated in yellow) were drawn across these cells [a (9 dpi), d (14 dpi), and g (21 dpi) and fluorescence (pixel) intensities along the transect lines were measured. Fluorescence intensity values of such transect lines were plotted along several points on the lines for each cell. A cell with one representative transect line and their corresponding intensity plot is shown here.

Together, these results demonstrate that Lb undergoes a developmentally regulated subcellular redistribution during nodule maturation.

### Rhizobial load increases with the progress of Nodulation

To determine whether the changes in the sub-cellular localization of Lb is correlated with rhizobial colonization. we examined bacterial abundance using propidium iodide (PI) staining. A progressive increase in rhizobial load was observed from early (9 dpi; Fig. 1A, b) to mature nodules (21 dpi; Fig. 1A, h). Consistent with this observation, quantitative real-time PCR analysis of the nitrogenase marker gene *nifH* showed a similar increasing trend during nodule development (Supplementary Fig. S1).

### Phosphorylated leghemoglobin emerges during mid-stage nodule development

Previous in vitro studies demonstrated that Lb undergoes serine phosphorylation at residues S45, resulting in reduced oxygen sequestration capacity (Bhar et al., 2015). Given that Lb is known to play a dual role in nodules, maintaining low free oxygen concentrations while simultaneously facilitating oxygen diffusion to respiring bacteroids (Appleby, 1984; Ott et al., 2005), we hypothesized that serine phosphorylation may represent a developmentally regulated mechanism to modulate Lb function during nodulation. To address this, FIHC was performed using phospho-peptide specific antibodies targeting S45P and S55P residue of Lb, along with antibodies recognizing their non-phosphorylated counterparts.

At 9 dpi, non-phosphorylated forms (LbS45NP and LbS55NP) were detected (Fig. 2A, b; Supplementary Fig. S2A, b respectively), consistent with total Lb accumulation at this stage (Fig. 1). In contrast, phosphorylated forms LbS45P and LbS55P were not detected until 14 dpi. At 14 dpi, phosphorylated Lb localized predominantly to the cell periphery (Fig. 2A, c; triangle, Supplementary Fig. S2A, c; triangle). By 21 dpi, phosphorylated forms increased substantially and were distributed throughout the cytoplasm (Fig.2A, e; star, Supplementary Fig. S2A, c; star).

**Figure 2.**
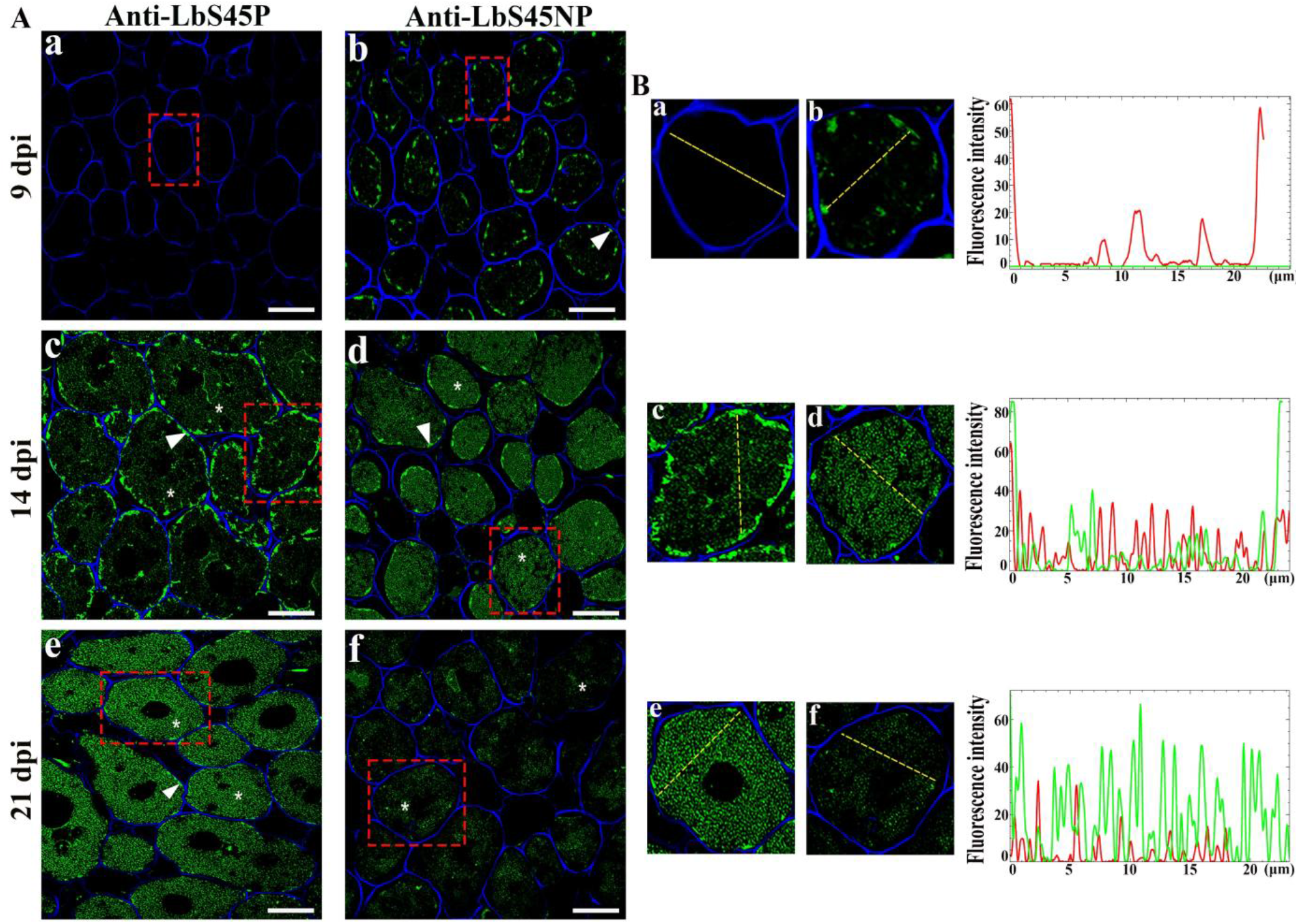
Spatio-temporal distribution of LbS45P and LbS45NP in *Lotus japonicus* nodules. **(A)** Nodules harvested at 9 dpi (a & b), 14 dpi (c & d) and 21 dpi (e & f) were processed for immunohistochemistry as mentioned earlier. LbS45P (green) localization is shown in panels a, c, and e and LbS45NP (green) localization is shown in panels b, d and f. _*_ Indicates cytoplasmic and ▲ demarcate cell periphery localization of Lb. Scale bars = 20 µm. Fluorescence intensity profiles of LbS45P (green) and LbS45NP (red) along the yellow transect lines in selected cells (a, c & e) and (b, d & f) respectively were determined and plotted as before. One such representative plot was shown for each sample.

Quantitative analyses of fluorescence intensities revealed a progressive increase in phosphorylated Lb (S45P and S55P) with nodule maturation, accompanied by a relative decline in non-phosphorylated forms (Fig. 2B; Supplementary Fig. S2B). Notably, S45 phosphorylation (Fig. 2B,) showed a stronger accumulation compared to S55 phosphorylation (Supplementary Fig. S2; B), suggesting potential site-specific regulatory differences.

### Lb phosphorylation first detectable at 11 dpi

To precisely determine the onset of Lb phosphorylation, nodules were analyzed at intermediate time points (10–13 dpi). Phosphorylated Lb became detectable at 11 dpi (Fig. 3A, c; triangle, Supplementary Fig. S3A, c; triangle), appearing as a peripheral signal adjacent to the cell wall.

**Figure 3.**
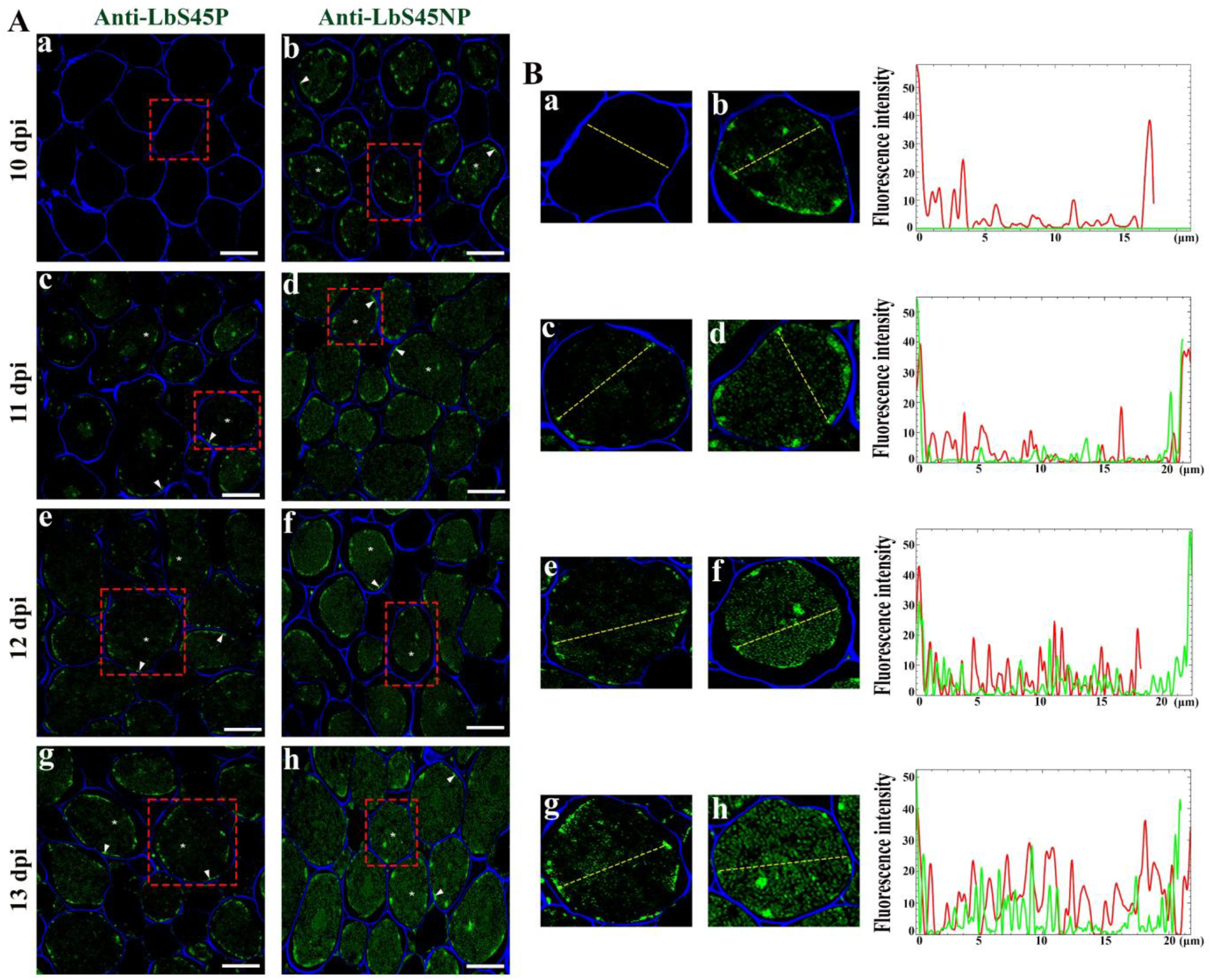
First evidence for spatial and temporal detection of LbS45 phosphorylation in *Lotus japonicus* nodules during nodulation. **(A)** Nodules harvested at 10 dpi (a & b), 11 dpi (c & d), 12 dpi (e & f), and 13 dpi (g & h) were processed for immunohistochemistry. LbS45P (green) localizations are shown in panel A (a, c, e, & g) and LbS45NP (green) localizations are shown in panel A (b, d, f & h). _*_ Indicates cytoplasmic and ▲demarcate cell periphery localization of Lb. Scale bars = 20 µm. **(B)** Fluorescence intensity profiles of LbS45P (green) and LbS45NP (red) along the yellow transect lines in selected cells (a, c, e & g) and (b, d, f & h) respectively were determined and plotted as before. One such representative plot was shown for each sample.

### Phosphorylated Lb initially localizes to the Plasma Membrane

To determine whether the peripheral localization of phosphorylated Lb corresponds to the plasma membrane, FIHC co-localization experiments were performed using phospho-specific antibodies (S45P and S55P) together with the plasma membrane marker LjPIP2 (Plasma Membrane Intrinsic Protein 2). At 14 dpi, phosphorylated Lb (S45P and S55P) exhibited clear co-localization with PIP2 (Fig. 4, c; triangle Supplementary Fig. S4, c; triangle), confirming its association with the plasma membrane.

**Figure 4.**
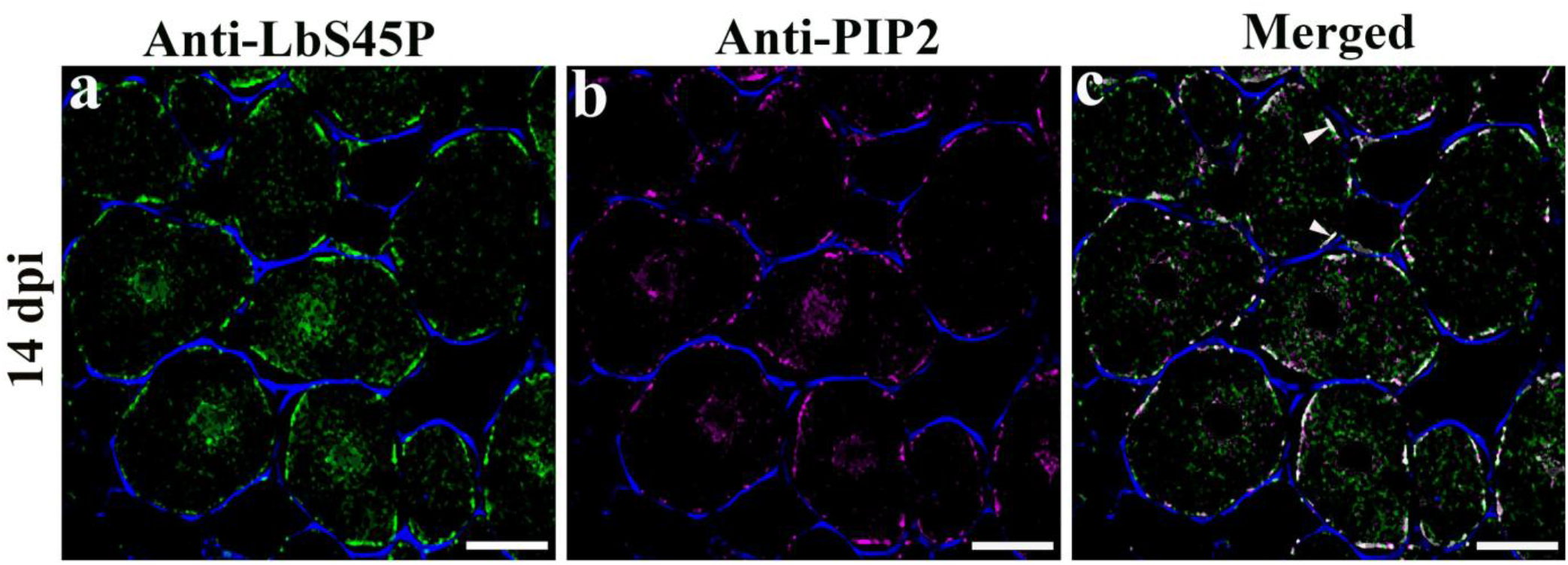
Phosphorylated Lb Localizes to the Plasma Membrane. Nodules harvested at 14 dpi (a, b & c) were processed for immunohistochemistry as mentioned earlier. LbS45P (green) localization is shown in panel a and PIP2 (magenta) localization is shown in panel b. Merged images are shown co-localization (off-white) in panel c (white triangle). Cell walls were visualized with Calcofluor White (blue) in all samples. Bars = 20 μm

These findings suggest that serine phosphorylation may promote transient membrane association of Lb prior to its redistribution into the cytoplasm during nodule maturation.

### Biochemical validation of developmentally regulated Lb phosphorylation

To validate the FIHC observations, total nodule proteins from 0, 9, 11, 14, and 21 dpi were subjected to immunoprecipitation using anti-Lb antibodies followed by immunoblotting with phospho- and non-phospho-specific peptide antibodies.

Non-phosphorylated Lb (S45NP and S55NP) was detectable at 9 dpi (Fig. 5A, b, L2; Supplementary Fig. S5A, b, L2 respectively), whereas phosphorylated forms (S45P and S55P) first appeared at 11 dpi (Fig. 5A, a, L3; Supplementary Fig. S5A, a, L3 respectively). Densitometric analysis revealed progressive accumulation of phosphorylated Lb took place through 21 dpi (Fig. 5B, green bars; Supplementary Fig. S5B, green bars). In contrast, non-phosphorylated forms increased up to 14 dpi but declined markedly at 21 dpi (Fig. 5B, red bars, Supplementary Fig. S5B, red bars).

**Figure 5.**
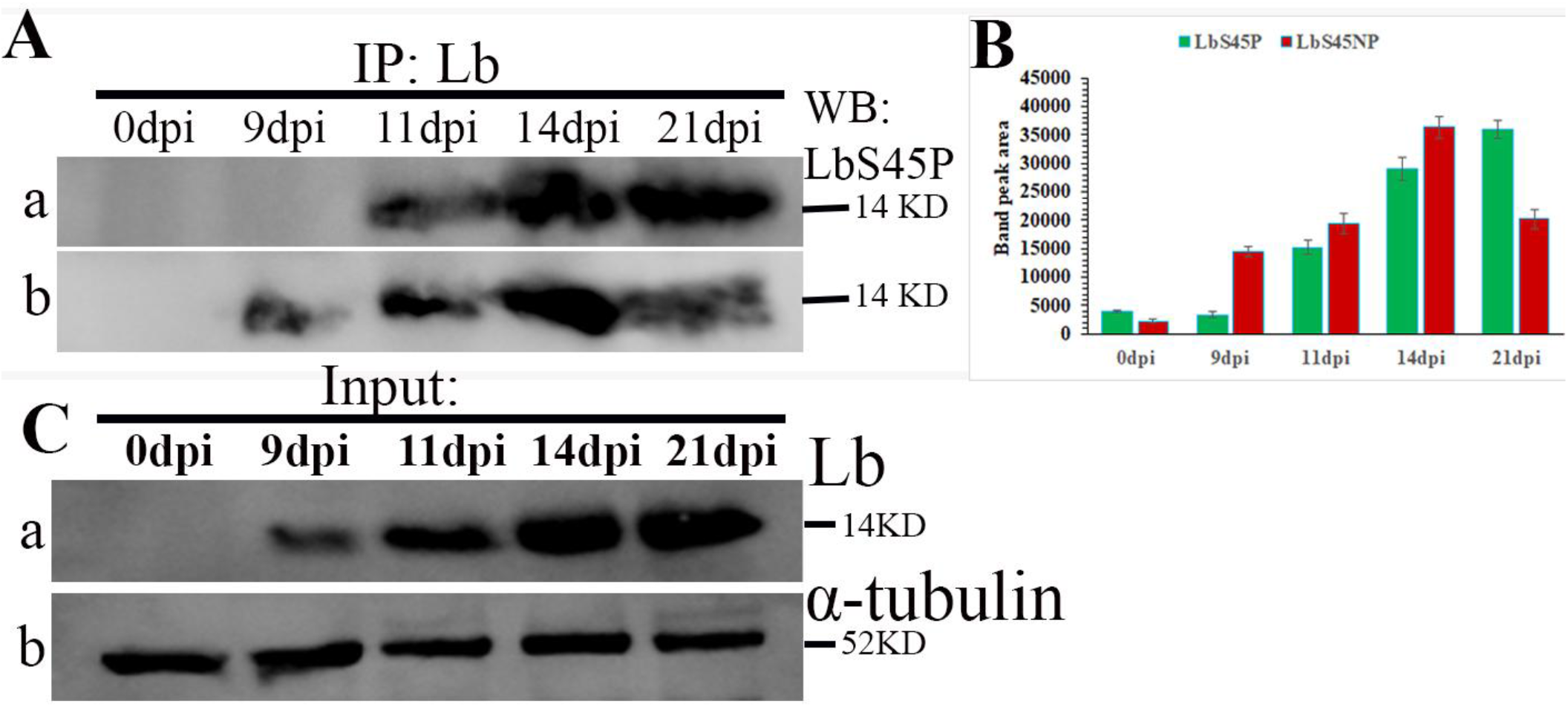
Developmental accumulation of LbS45P and LbS45NP in nodule lysates at different dpi during nodulation. Total nodule protein (80 μg), prepared using a denaturing extraction method (Methods), was subjected to immunoprecipitation with anti-Lb antibodies and subsequent immunoblotting. **(A)** Immunoprecipitated samples from nodules at 0, 9, 11, 14, and 21 days post-inoculation (dpi) (lanes 1–5) were immunoblotted with anti-LbS45P (Fig. 5A, a) and anti-LbS45NP (Fig. 5A, b) antibodies to detect S45-phosphorylated and non-phosphorylated Lb (∼14 kDa), respectively. **(B)** Bar graphs show densitometric quantification of LbS45P (Fig. 5B, green bars) and LbS45NP (Fig. 5B, red bars) band intensities (∼14 kDa), measured using ImageJ across different dpi. **(C)** Input controls showing total Lb (Fig. 5C, a) levels and α-tubulin (∼52 kDa) (Fig. 5C, b) as a loading control across developmental stages.

These biochemical data corroborate the temporal patterns observed by FIHC.

## Discussion

Nitrogen fixation is a highly energy-intensive process requiring approximately 16 ATP molecules to reduce one molecule of N_2_ to NH_3_ (Appleby, 1984; Udvardi and Poole, 2013). This high energy demand is met by aerobic respiration of rhizobia; however, the nitrogenase enzyme responsible for this reaction is extremely sensitive to oxygen. This inherent conflict between oxygen-dependent respiration and oxygen-sensitive nitrogenase activity constitutes the well-known oxygen paradox of symbiotic nitrogen fixation (Appleby et al., 1984).

In indeterminate nodules, this paradox is partially resolved through zonal organization that enables differential oxygen regulation (Kawashima et al., 2001). In contrast, determinate nodules such as those of *Lotus japonicus* lack such spatial differentiation (Udvardi and Poole, 2013), suggesting that intracellular regulatory mechanisms play a central role in maintaining oxygen homeostasis during nodule development. Post-translational modifications, particularly phosphorylation, represent a plausible mechanism for such regulation.

Previous studies, results from our and other laboratories showed that *in vitro* phosphorylation of Lb from *Lotus japonicus* at specific serine residues (S45 and S55) significantly reduces its oxygen sequestration capacity by altering the heme-binding pocket (Bhar et al., 2015). Subsequent phosphoproteomic analyses identified phosphorylation of Lb at a homologous site in *Medicago truncatula*, suggesting that this modification is evolutionarily conserved and may play a regulatory role during symbiotic nitrogen fixation (Marx et al., 2016). However, the spatial and developmental relevance of Lb phosphorylation in vivo remained unclear.

In the present study, we examined the spatio-temporal dynamics of Lb (fig. 1) and its phosphorylation at S45 and S55 residue during nodule development to understand how this modification linked to changes in its subcellular localization within infected nodule cells. Our results show that serine phosphorylation of Lb initially becomes detectable in the plasma membrane around 11dpi (fig. 3A, c) and subsequently redistributed into the cytoplasm as nodules mature (fig. 2A, e). This transition coincides with increasing rhizobial colonization and metabolic demand.

Previous work from our laboratory demonstrated that Lb interacts with another late nodulin protein Nlj16 leading to enhanced oxygen sequestration capacity of Lb (Ghosh et al., 2019). Interestingly, recent work from our laboratory also revealed that Lb is recruited to the cell membrane through interaction with Nlj16 (Ghosh et al., 2022). Moreover, our preliminary data indicates that phosphorylation of Lb disrupts its interaction with Nlj16. Thus, the fact that the association of Lb with the cell membrane via Nlj16 is reduced upon subsequent serine phosphorylation of Lb resulting in its redistribution into the cytoplasm. This interpretation is consistent with the observed spatial dynamics of phosphorylated Lb during nodule maturation. Our observations support a developmental model in which Lb function shifts from oxygen sequestration to controlled oxygen delivery during nodule maturation (Fig. 6). During early stages of nodulation, when rhizobial populations are relatively low, non-phosphorylated Lb interacts with Nlj16 at the plasma membrane and exhibits high oxygen affinity, effectively sequestering oxygen and protecting nitrogenase activity. As nodules mature and bacteroid numbers increase, respiratory demand rises substantially. Under these conditions, excessive oxygen sequestration could limit oxygen availability for bacteroid respiration. Phosphorylation of Lb at S45 and S55 likely reduces its oxygen-binding affinity and disrupts its interaction with Nlj16, resulting in release from the membrane and redistribution throughout the cytoplasm. This transition may facilitate localized oxygen release near symbiosomes (Appleby, 1984; Udvardi and Poole, 2013), supporting aerobic respiration and ATP production while maintaining oxygen levels below nitrogenase-inhibiting thresholds.

**Fig. 6.**
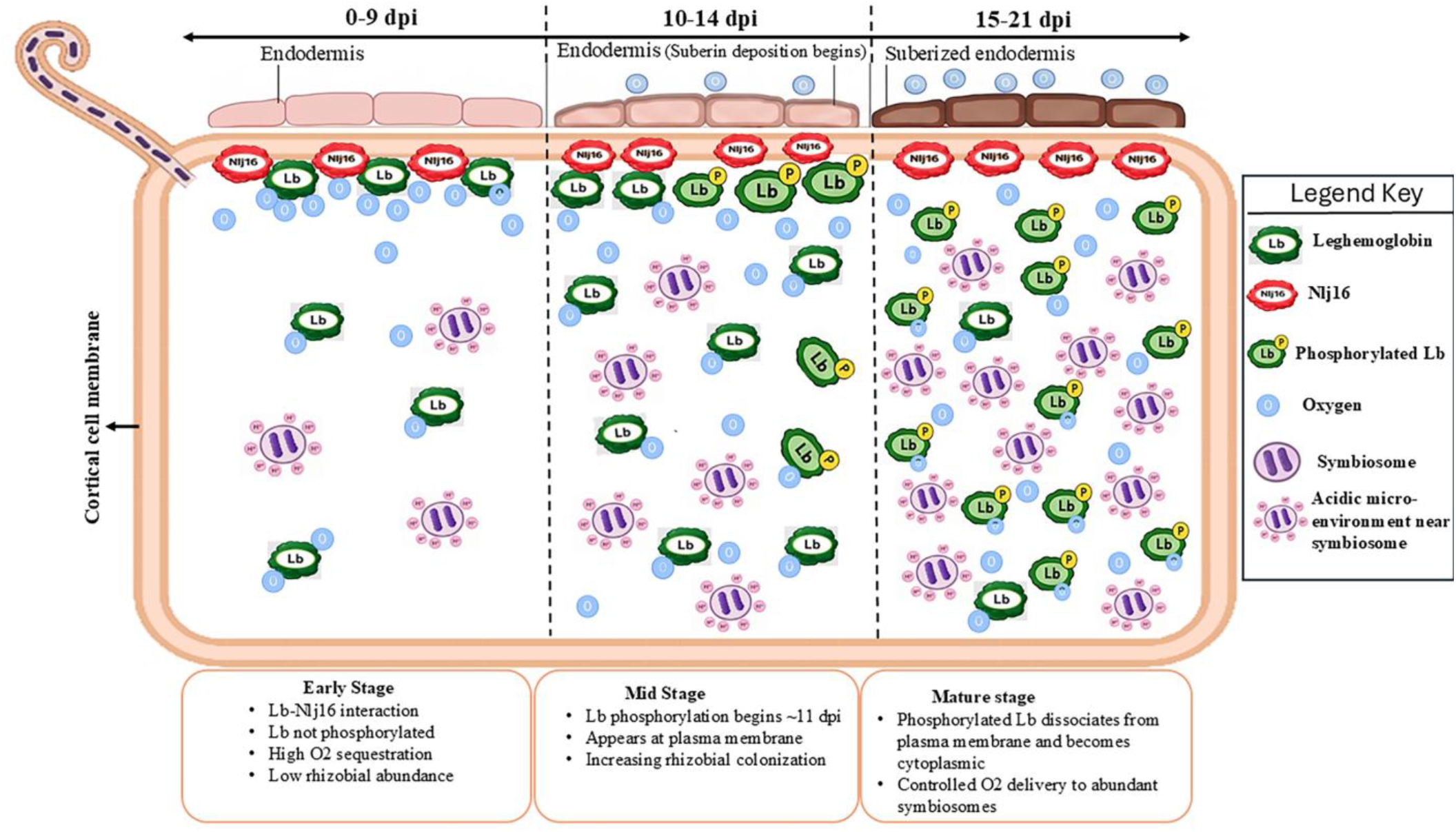
Developmental regulation of leghemoglobin (Lb) phosphorylation during nodule maturation. Schematic model illustrating the spatiotemporal dynamics of Lb localization and phosphorylation in *Lotus japonicus* nodules. During early stages (0–9 dpi), non-phosphorylated Lb associates with Nlj16 at the plasma membrane, promoting efficient oxygen sequestration under low rhizobial load. At the onset of nodule maturation (10–14 dpi), Lb phosphorylation begins (∼11 dpi) and is primarily detected at the plasma membrane. In mature nodules (15–21 dpi), phosphorylated Lb dissociates from the membrane and redistributes to the cytoplasm, enabling controlled oxygen delivery to symbiosomes. This transition supports increased bacteroid respiration while maintaining oxygen concentrations below nitrogenase-inhibiting level

Additional factors may further reinforce this regulatory mechanism. Previous studies have shown that the acidic microenvironment surrounding the peribacteroid membrane promotes oxygen release from Lb (Pierre et al., 2013). Thus, phosphorylation-dependent reduction in oxygen-binding affinity, together with oxygen gradients and local pH effects, may synergistically enhance oxygen availability for bacteroid respiration while maintaining concentrations below nitrogenase-inhibiting levels. Moreover, structural adaptations such as suberin deposition in the endodermal tissues of later dpi nodules may contribute to restricting oxygen diffusion into the nodule interior (Venado et al., 2022).

Importantly, our findings establish a strong correlation between phosphorylation, subcellular localization, and nodule maturation. However, whether phosphorylation directly drives these changes remains to be determined. Future studies using phospho-mutant and phospho-mimic variants of Lb will be necessary to establish causal relationships.

Together, these findings suggest that developmentally regulated specific serine phosphorylation of Lb coordinates its localization, protein interactions, and oxygen-binding properties to dynamically regulate oxygen availability during symbiotic nitrogen fixation in determinate nodules. This mechanism provides an additional layer of control over oxygen homeostasis and highlights phosphorylation-dependent modulation of leghemoglobin as a key regulatory strategy supporting efficient nitrogen fixation.

## Acknowledgements

This work is funded by Grants from Govt. of India DST SERB (EMR/2017/004234) and DST ANRF (CRG/2023/005319) to AS. MAA and AD were supported by PhD research fellowships from the University Grants Commission and Council of Scientific and Industrial Research, India. This research is also supported by DST-FIST (Department of Biochemistry), DBT-Builder and DST-PURSE programs of University of Calcutta, India.

## Author Contribution

MAA and AS conceived and designed the study. MAA and AD performed the experiments. MAA and AS analysed and interpreted the data. MAA, AD, and AS wrote the manuscript. AS supervised the project and secured funding.

## Supplementary Data

**Supplementary Figure S1**. Rhizobial load with the progress of Nodulation.

**Supplementary Figure S2**. Spatio-temporal distribution of LbS55P and LbS55NP in *Lotus japonicus* nodules.

**Supplementary Figure S3**. First evidence for spatial and temporal detection of LbS55 phosphorylation in *Lotus japonicus* nodules during nodulation.

**Supplementary Figure S4**. Phosphorylated Lb Localizes to the Plasma Membrane.

**Supplementary Figure S5**. Developmental accumulation of LbS55P and LbS55NP in nodule lysates at different dpi during nodulation.

## Supplementary Information for

**Supplementary figure S1:**
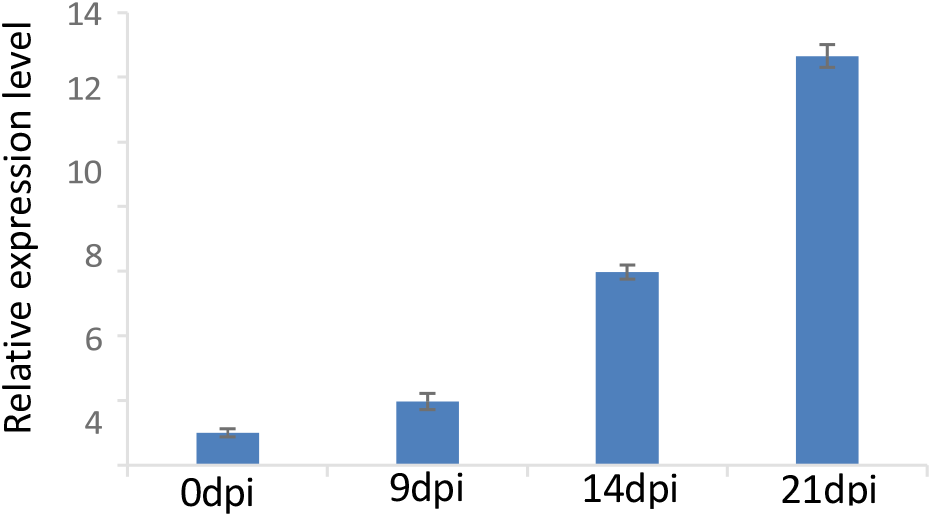
Rhizobial load with the progress of Nodulation. Relative mRNA expression levels of nifH in the nodules of *Lotus japonicus* harvested at 0 dpi,9 dpi,14 dpi and 21 dpi were determined using real time PCR experiments. 2^-ΔΔCt^ values were calculated using sigA as the internal control. Error bars represent the standard deviations of the three independent experiments.

**Supplementary Figure S2:**
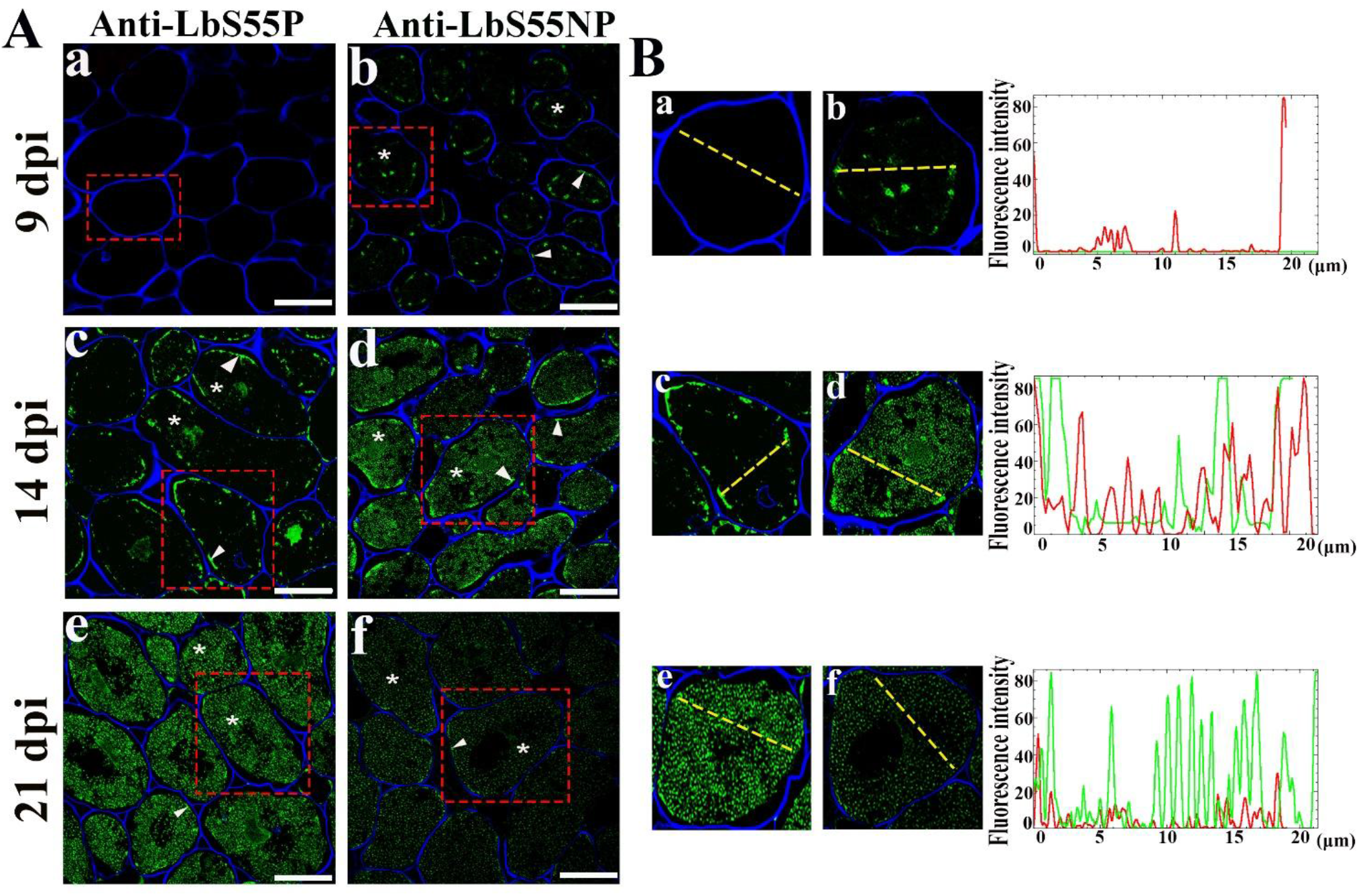
Spatio-temporal distribution of LbS55P and LbS55NP in *Lotus japonicus* nodules. **(A)** Nodules harvested at 9 dpi (a & b), 14 dpi (c & d) and 21 dpi (e & f) were processed for immunohistochemistry as mentioned earlier. LbS55P (green) localization is shown in panels a, c, and e and LbS55NP (green) localization is shown in panels b, d and f. * Indicates cytoplasmic and ▲ demarcate cell periphery localization of Lb. Scale bars = 20 μm. **(B)** Fluorescence intensity profiles of LbS55P and LbS55NP along the yellow transect lines in selected cells (a, c & e) and (b, d & f) respectively were determined and plotted as before. One such

**Supplementary figure S3:**
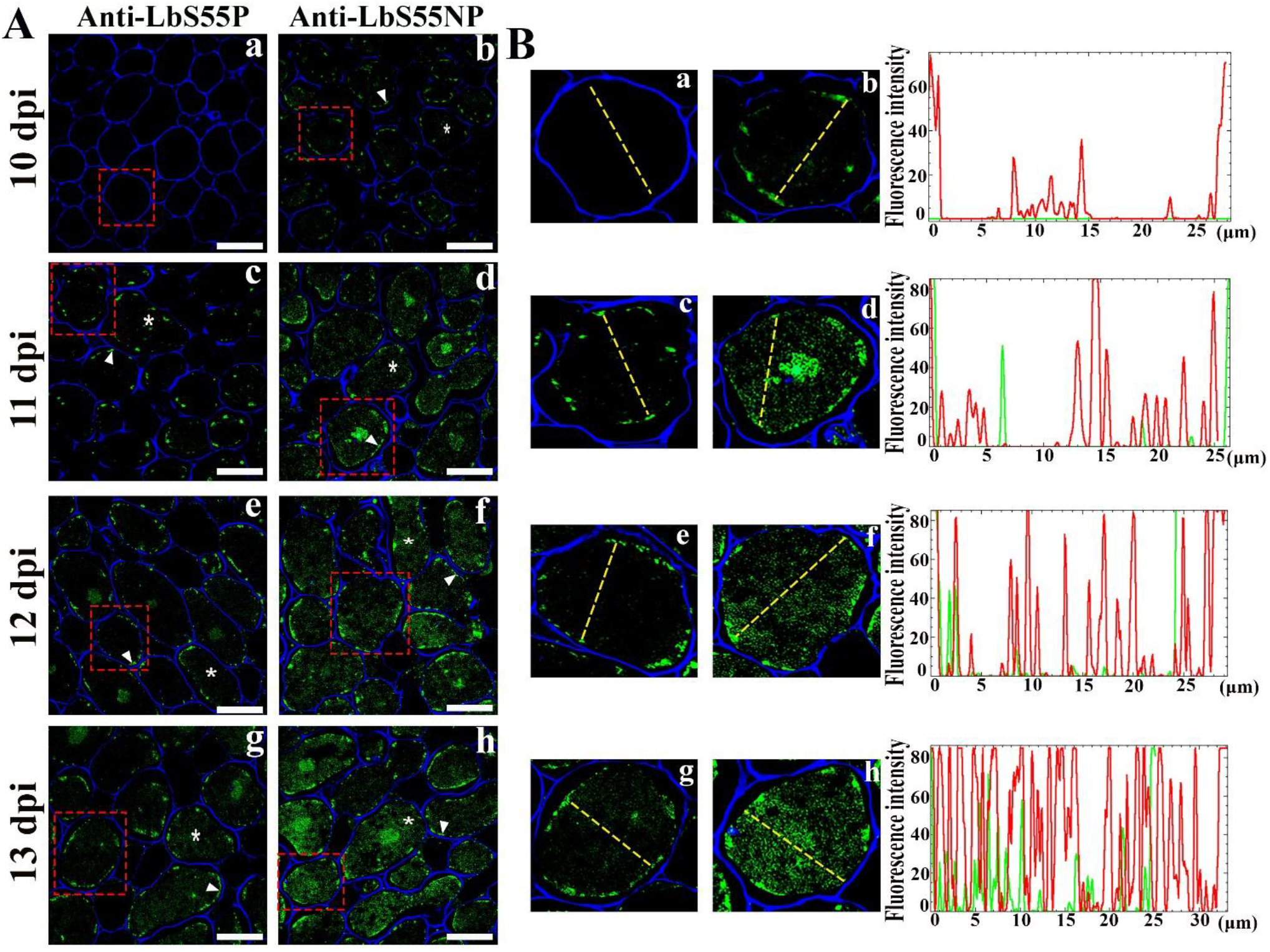
First evidence for spatial and temporal detection of LbS55 phosphorylation in *Lotus japonicus* nodules during nodulation. (A) Nodules harvested at 10 dpi (a & b), 11 dpi (c & d), 12 dpi (e & f), and 13 dpi (g & h) were processed for immunohistochemistry. LbS55P(green) localizations are shown in panel A (a, c, e, & g) and LbS45NP (green) localizations are shown in panel A (b, d, f & h). * Indicates cytoplasmic and ▲ demarcate cell periphery localization of Lb. Scale bars = 20 μm. **(B)** Fluorescence intensity profiles of LbS45P (green) and LbS45NP (red) along the yellow transect lines in selected cells (a, c, e & g) and (b, d, f & h) respectively were determined and plotted as before. One such representative plot was shown for each sample.

**Supplementary figure S4:**
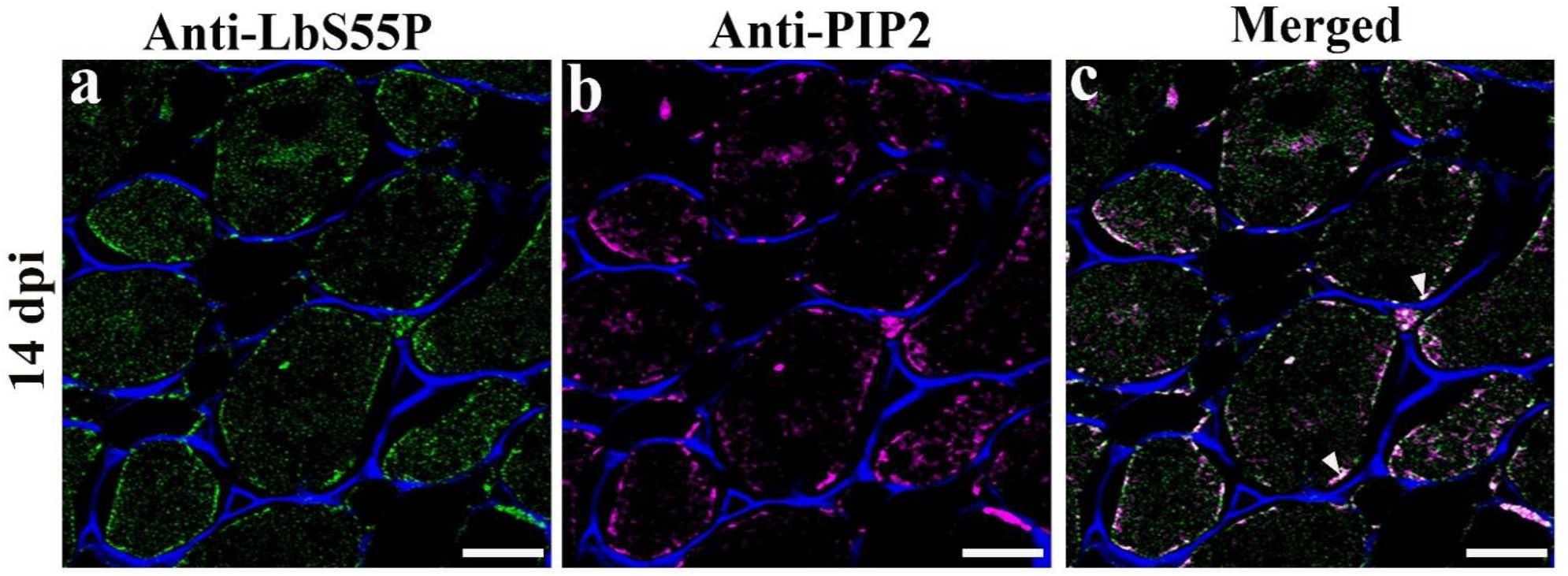
Phosphorylated Lb Localizes to the Plasma Membrane. Nodules harvested at 14 dpi (a, b & c) were processed for immunohistochemistry as mentioned earlier. LbS55P (green) localization is shown in panel a and PIP2 (magenta) localization is shown in panel b. Merged images are shown co-localization (off-white) in panel c (white triange). Cell walls were visualized with Calcofluor White (blue) in all samples. Bars = 20 μm.

**Supplementary Figure S5:**
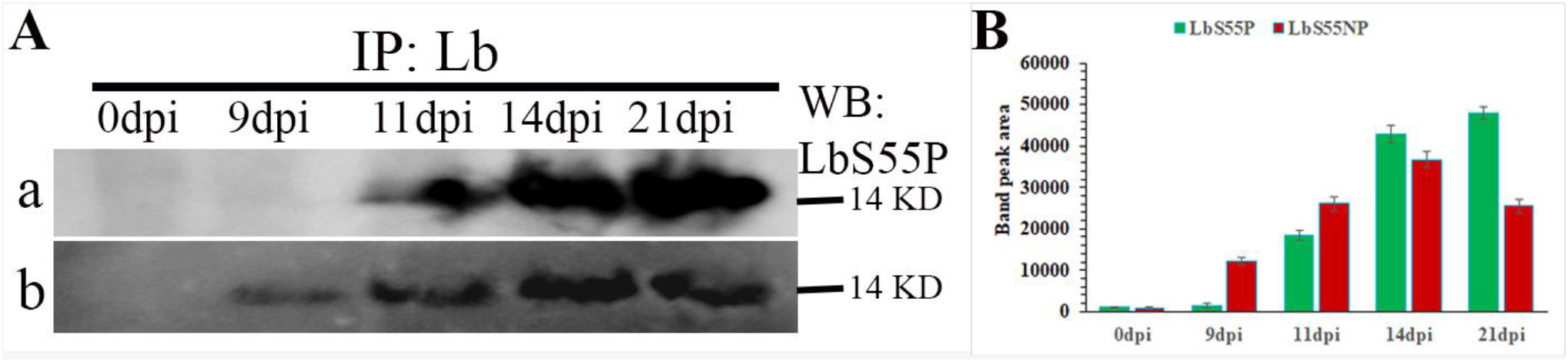
Developmental accumulation of LbS55P and LbS55NP in nodule lysates at different dpi during nodulation. Total nodule protein (80 μg), prepared using a denaturing extraction method (Methods), was subjected to immunoprecipitation with anti-Lb antibodies and subsequent immunoblotting. **(A)** Immunoprecipitated samples from nodules at 0, 9, 11, 14, and 21 days post-inoculation (dpi) (lanes 1 −5) were immunoblotted with anti-LbS55P (Fig. S5A, a) and anti-LbS55NP (Fig. S5A, b) antibodies to detect S55-phosphorylated and non-phosphorylated Lb (∼14 kDa), respectively. **(B)** Bar graphs show densitometric quantification of LbS55P (Fig. S5B, green bars) and LbS55NP (Fig. S5B, red bars) band intensities (∼14 kDa), measured using ImageJ across different dpi.

